# Multiscale Physical Effects of CpG Methylation on DNA Mechanics, Nucleosome Wrapping, and Chromatin Condensates

**DOI:** 10.64898/2026.04.13.715649

**Authors:** LJC Daris, A Rizvi, A Singh, J Hutchings, LA Lamers, EA Montabana, D Kimanius, PR Banerjee, JM Eeftens, S Sanulli

**Author notes:** equal contributions (LJC Daris, A Rizvi). Co-corresponding authors Correspondence to Jorine M Eeftens and Serena Sanulli.

## Abstract

DNA methylation at CpG dinucleotides is a central epigenetic modification linked to transcriptional repression and heterochromatin formation, yet its physical impact on chromatin organization remains incompletely understood. Here, we integrate single-molecule force spectroscopy, FRAP, and nanorheology to quantify how CpG methylation alters chromatin mechanics across scales. Optical tweezer-based measurements show that CpG methylation modestly increases DNA contour and persistence lengths, indicating a slightly extended and stiffer DNA helix. When assembled into nucleosomes, CpG methylation weakens both outer-and inner turn wrapping, reducing unwrapping forces and per-nucleosome unwrapping energy. On a higher-order assembly scale, although CpG methylated and unmethylated chromatin fibers display similar thresholds for salt-induced phase separation, methylated condensates exhibit markedly reduced internal mobility: FRAP reveals slower chromatin exchange, nanorheology shows increased viscoelasticity and longer relaxation times. Finally, CpG methylation enhances HP1α–chromatin interactions, lowering the HP1α concentration required for condensation, despite no change in HP1α binding to naked DNA. Together, these results show that CpG methylation remodels chromatin mechanics by tuning DNA rigidity, nucleosome wrapping energetics, and condensate material properties, providing a multiscale physical framework for how DNA methylation contributes to heterochromatin.

## Introduction

The eukaryotic genome is packaged within the confined volume of the nucleus as chromatin, forming hierarchically organized structures that span multiple length scales^1,2^. The first level of chromatin organization involves the wrapping of DNA around an octamer of histone proteins to form nucleosomes, the basic repeating units of chromatin^3,4^. Arrays of nucleosomes fold further into higher-order structures, including chromatin fibers, topologically associated domains, and chromosome-scale compartments^1,2^. Beyond enabling efficient genome packaging, chromatin folding functionally organizes the genome to regulate genome processes, such as transcription, replication, and gene silencing^5,6^.

Chromatin exists along a continuum of compaction states that broadly correlate with transcriptional states^7–10^. Euchromatin, which is relatively open and accessible, is associated with active transcription, permissive histone modifications, and dynamic chromatin-binding factors^11^. In contrast, heterochromatin represents a more compact, transcriptionally repressive state that plays essential roles in genome stability, developmental gene silencing, and nuclear organization^11,12^. Heterochromatin domains can form constitutively at repetitive and pericentromeric regions, or facultatively in a cell-type–specific and developmentally regulated manner^13–15^. These compact domains not only restrict transcriptional access but also serve as structural scaffolds involved in nuclear organization^12,16,17^.

At the molecular level, heterochromatin is defined by specific biochemical hallmarks. Among the most conserved and functionally linked are histone H3 lysine 9 trimethylation (H3K9me3) and DNA methylation^12,18–20^. H3K9me3 is recognized by heterochromatin protein 1α (HP1α), which binds nucleosomes, promotes chromatin compaction in condensates, and drives the formation of microscopically visible heterochromatic foci^12,18,21–23^. DNA methylation at CpG dinucleotides similarly contributes to gene repression by recruiting methyl-CpG–binding proteins and co-repressor complexes, and by sterically hindering the binding of transcriptional activators^19,24,25^. Together, these marks form mutually reinforcing pathways that stabilize silent chromatin domains and preserve cell identity^19^.

In DNA methylation, a methyl group is covalently added to the 5-carbon position of the cytosine ring (5mC), predominantly at CpG dinucleotides in mammalian genomes^25^. This modification serves as a fundamental epigenetic signal that regulates genome function by recruiting methyl-binding proteins or by directly modulating DNA–protein interactions^25–27^. In living cells, DNA methylation increases chromatin condensation and decreases overall DNA flexibility, even in the absence of the dedicated reader machinery, indicating that beyond serving as a docking site for chromatin factors, it can intrinsically influence the physical properties of DNA and chromatin^28^. However, the mechanism through which DNA methylation affects the mechanics and organization of chromatin remains controversial. Computational and biophysical studies have suggested that methylation reduces DNA flexibility and disfavors nucleosome assembly, whereas other *in vitro* studies report enhanced histone–DNA affinity and increased nucleosome compaction^29–33^. These discrepancies likely reflect the complexity of chromatin and the context-dependent nature of nucleosome stability within higher-order structures.

Here, we address this gap by combining single-molecule force spectroscopy, Fluorescence Recovery After Photobleaching (FRAP), and nanorheology to interrogate the effects of CpG methylation from the level of naked DNA to reconstituted nucleosomes and higher-order chromatin condenstes. This integrated approach enables direct comparison of DNA methylation’s mechanical and structural impact across scales and provides a physical framework for understanding how this epigenetic modification shapes chromatin architecture and dynamics.

## Results

### CpG Methylation Affects Mechanical Properties of DNA

Optical tweezers enable direct mechanical characterization of DNA by tethering single DNA molecules between two optically-trapped beads and applying controlled stretching forces. To investigate the effect of cytosine methylation on the mechanical properties of DNA, a ∼6.8 kb plasmid containing 12 repeats of the Widom 601 nucleosome positioning sequence was linearized, and the 5’-overhangs were biotinylated to allow tethering to trapped beads in our optical tweezers. The CpG methyltransferase (MTase) M.SssI was used to methylate all the 482 cytosines at the C-5 position present within the double-stranded CG dinucleotide sequence (**Fig. 1A**). Digestion with the CpG methylation-sensitive restriction enzyme HpaII confirmed that, in presence of the MTase and methyl donor S-adenosylmethionine (SAM), the DNA was fully methylated and protected against degradation (**Fig. S1**).

**Figure 1.**
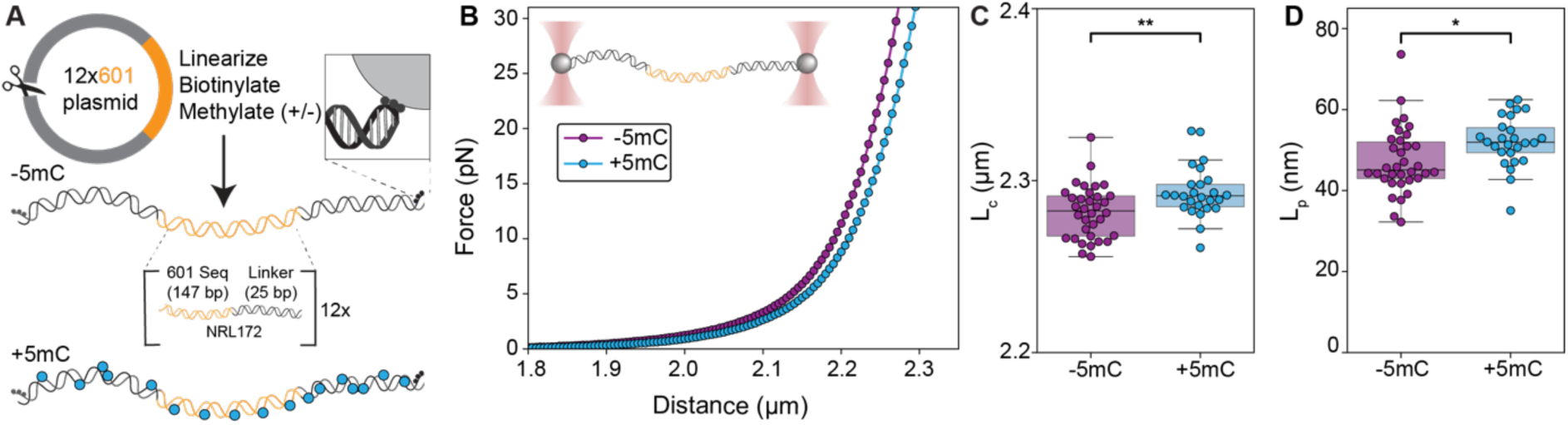
CpG methylation alters contour length (L*_C_*) and persistence length (L*_P_*) of dsDNA. **A)** Schematic of DNA construct preparation for single-molecule force spectroscopy. A ∼6.8 kb plasmid was linearized with the restriction enzyme NotI and the 5’-overhangs were biotinylated using Klenow fragment. The CpG methyltransferase (MTase) M.SssI was used to methylate all cytosines at the C-5 position (5mC) within the double-stranded CG dinucleotide sequence. **B)** Interpolation of force-extension data shows that the averaged force-distance (F,d) curves for unmethylated (−5mC) and methylated (+5mC) DNA follow different trajectories, reflecting variation in the mechanical properties upon CpG methylation. **C**) CpG methylation significantly increases the DNA contour length (L*_C_*) (Student’s t-test, p < 0.001, n = 34 and n = 26 for -5mC and +5mC, respectively). Box plots show quartiles; whiskers show the distribution excluding outliers. **D**) CpG methylation significantly increases the DNA persistence length (L*_p_*) from 47 ± 8 nm (unmethylated) to 52 ± 7 nm (methylated) (Student’s T-test, p < 0.05, n = 34 and n = 26 for -5mC and +5mC, respectively).

**Figure 1.**
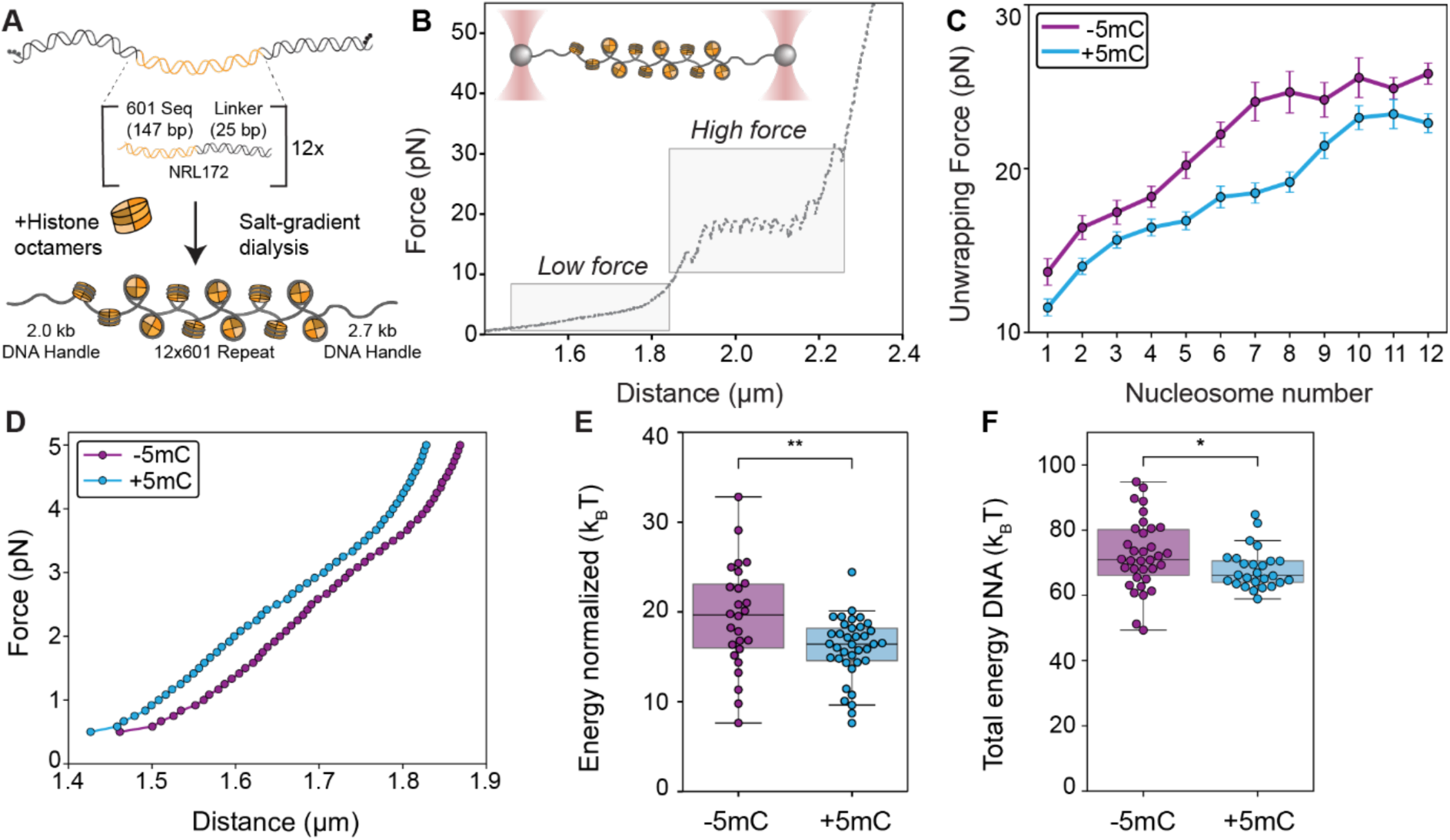
CpG methylation weakens nucleosome stability and lowers the energy required to stretch chromatin fibers. **A)** Schematic of chromatin reconstitution. 12×601 DNA template and recombinant human histone octamers were assembled into nucleosomal arrays by salt-gradient dialysis. Gradual reduction of salt promotes spontaneous nucleosome formation at each 601-positioning sequence. **B**) Representative F,d-curve of a 12-mer nucleosomal array. Chromatin fibers were tethered between optically trapped streptavidin-coated polystyrene beads and stretched by gradual displacement of one trap. The force response of chromatin presents two regimes: a low–force regime attributed to intra-nucleosome interaction unstacking and outer turn DNA unwrapping, and a high–force regime corresponding to unwrapping of the nucleosome inner turns. **C**) The force required to unwrap the nucleosome inner DNA turn is significantly lower in +5mC arrays, indicating that DNA methylation destabilizes the inner turn (Student’s t-test for nucleosomes 1, 2, 5, 7, 8, 9 and 12. Mann-Whitney U-test for nucleosomes 3, 4, 6, 10 and 11. p < 0.001 for 6-8; p < 0.01 for 5; p < 0.05 for 1,2 and 4; ns for 3 and 9-12). Error bars represent SEM. **D**) Force trace data was interpolated to obtain average F,d-curves for stretching -5mC and +5mC chromatin fibers in the low force regime (0.5 pN < F < 5 pN). Similarity in the trajectories imply that unfolding of the compacted chromatin fiber is comparable for both conditions. **E**) Normalizing total energy by the number of nucleosomes per fiber reveals a significant decrease for methylated arrays (Student’s t-test: p < 0.01, n = 37 and n = 26 for +5mC and -5mC, respectively). **F**) The energy required to stretch the DNA in the same regime is significantly reduced for +5mC arrays (Mann-Whitney U-test: p < 0.05, n = 34 and n = 26 for +5mC and -5mC, respectively).

Single-molecule force spectroscopy was conducted by tethering the linear biotinylated DNA construct between two streptavidin-coated polystyrene beads that were optically trapped in a dual optical trap setup. Displacement of one of the traps was performed with a constant velocity of 20 nm/s. The elastic response of the tethered DNA induces forces on the stationary bead that are registered using back focal plane detection of scattered optical trapping light. The averaged Force,distance (F,d)-curves showed qualitative differences in the trajectories between methylated (+5mC) – and unmethylated (−5mC) DNA (**Fig. 1B**).

To relate these differences in elastic response to the mechanical properties of DNA, Odijk’s extensible worm-like chain (eWLC) model was used^34^. Fitting the force trace data to this model revealed that CpG methylation caused a modest but significant increase in the contour length (L*_c_*) of DNA (**Fig. 1C**), suggesting a slight increase in the spacing between base pairs. Additionally, the persistence length (L*_p_*), which reflects DNA structural flexibility, also increased upon methylation, indicating that +5mC DNA is less flexible than -5mC DNA (**Fig. 1D**). These results are consistent with previous reports that DNA methylation enhances DNA stiffness^29,35–39^.

Together, these data suggest that CpG methylation slightly extends the DNA helix while reducing its structural flexibility.

### CpG Methylation Decreases Stability of the Nucleosome Inner Turn and Alters DNA Wrapping

To determine the effect of 5mC on chromatin fibers, we reconstituted nucleosomal arrays using the same 6.8 kb DNA template as described above, containing 12 Widom 601 sequences spaced by 25 bp linkers. This results in a nucleosome repeat length (NRL) of 172 bp. Chromatin fibers were generated by salt dialysis of the template DNA with recombinant human histone octamers (**Fig. 2A**). Saturation of the 601 Widom sequences was confirmed by an electrophoretic mobility shift assay (EMSA) (**Fig. S2A, B**).

Single-molecule force spectroscopy was used to probe the mechanical stability of reconstituted chromatin fibers, following the same approach applied to naked DNA. Differently from the DNA, chromatin showed F,d-curves characterized by two distinct regimes that reflect different chromatin structural states **(Fig. 2B)**. In the low force regime (F < 10 pN), the applied force induces unwrapping of the outer DNA turns from nucleosomes and the disruption of any nucleosome-nucleosome interactions within the fiber^40^. Because chromatin fibers assembled with 10n+5 linker DNA lengths do not support intra-fiber nucleosome stacking^41,42^ the observed response in this regime reflects nucleosome unwrapping rather than higher-order folding transitions. In the high force regime (F > 10 pN), chromatin fibers are decompacted into a ‘beads-on-a-string’ conformation. Further increases in the applied force led to the unwrapping of ∼76 bp of inner turn DNA from each nucleosome, observed as a discrete drop in the recorded force accompanied by an increase in fiber extension. Each step corresponds to the unwrapping of an individual nucleosome, allowing estimation of nucleosome number per fiber. Consistent with a previous dynamic force spectroscopy model^43^, the unwrapping force for both +5mC and -5mC fibers increased progressively with each event, reflecting the non-cooperative nature of inner turn unwrapping (**Fig. 2C**)^44^. Rupture forces associated with inner turn unwrapping were quantified by identifying the peak force preceding each rupture event (**Fig. S2C**), informing on the resistance to mechanical disruption of the interactions of the DNA with the histone octamer. While both +5mC and -5mC chromatin fibers present a consistent increase of unwrapping forces for each successive unwrapping event, +5mC fibers consistently showed significantly lower unwrapping forces, indicating that 5mC destabilizes the nucleosome inner turn under applied force (**Fig. 2C**).

Since intra-fiber nucleosome stacking interactions are negligible in NRL172 chromatin fibers, the contribution of nucleosomes to the force response of chromatin in the low force regime is dominated by unwrapping of the nucleosomal DNA outer turns. The similar shape of the averaged F,d-curves illustrates that +5mC and -5mC chromatin fibers respond comparably under applied force (**Fig. 2D**). The leftward shift of the average curve for +5mC chromatin arises from the increased number of nucleosomes per fiber under this condition, which decreases the contour length L*_c_* of the fully wrapped chromatin fiber **(Fig. S2D)**. Because stretching in the low force regime is reversible^43,45^, the energy required to unwrap the nucleosomal outer turn can be calculated from the total area under the curve (AUC) and normalized to the number of nucleosomes per fiber (**Fig. S2D, E**). This analysis revealed that the unwrapping energy per nucleosome is lower for +5mC chromatin (**Fig. 2E**), indicating reduced nucleosome stability. This energy reflects contributions from both the unwrapping of the nucleosomal outer turn and the intrinsic stretching behavior of the underlying DNA. To assess the latter contribution, we quantified the energy required to stretch naked dsDNA (i.e. without nucleosomes) and found that 5mC significantly decreases the energy needed to extend DNA within this same regime (**Fig. 2F**). These results indicate that both reduced nucleosome outer turn stability and DNA contribute to the mechanical behavior of methylated chromatin fibers. The observed differences in unwrapping forces between methylated and unmethylated fibers suggest that CpG methylation affects how tightly DNA is wrapped around the histone octamer.

Together, our single-molecule analyses reveal that CpG methylation modulates chromatin mechanics by altering how DNA wraps around the histone octamer, weakening outer turn stability and subtly changing nucleosome geometry. Overall, CpG methylation reshapes chromatin properties by coupling changes in DNA mechanics to nucleosome stability.

### CpG Methylation Decreases Chromatin Dynamics within Condensates

To determine how methylation influences higher-order chromatin organization, we next examined the phase separation behavior of methylated and unmethylated fibers. Chromatin phase separation reflects multivalent inter-nucleosome interactions that drive formation of condensates, providing a direct readout of how DNA methylation affects these contacts^23,41^. Under physiological salt conditions, both –5mC and +5mC chromatin fibers underwent phase separation at comparable threshold concentration (c_phase_ ≈ 2.5 nM), indicating that CpG methylation does not substantially alter nucleosome-nucleosome interactions (**Fig. 3A, B; Fig. S3**).

**Figure 3.**
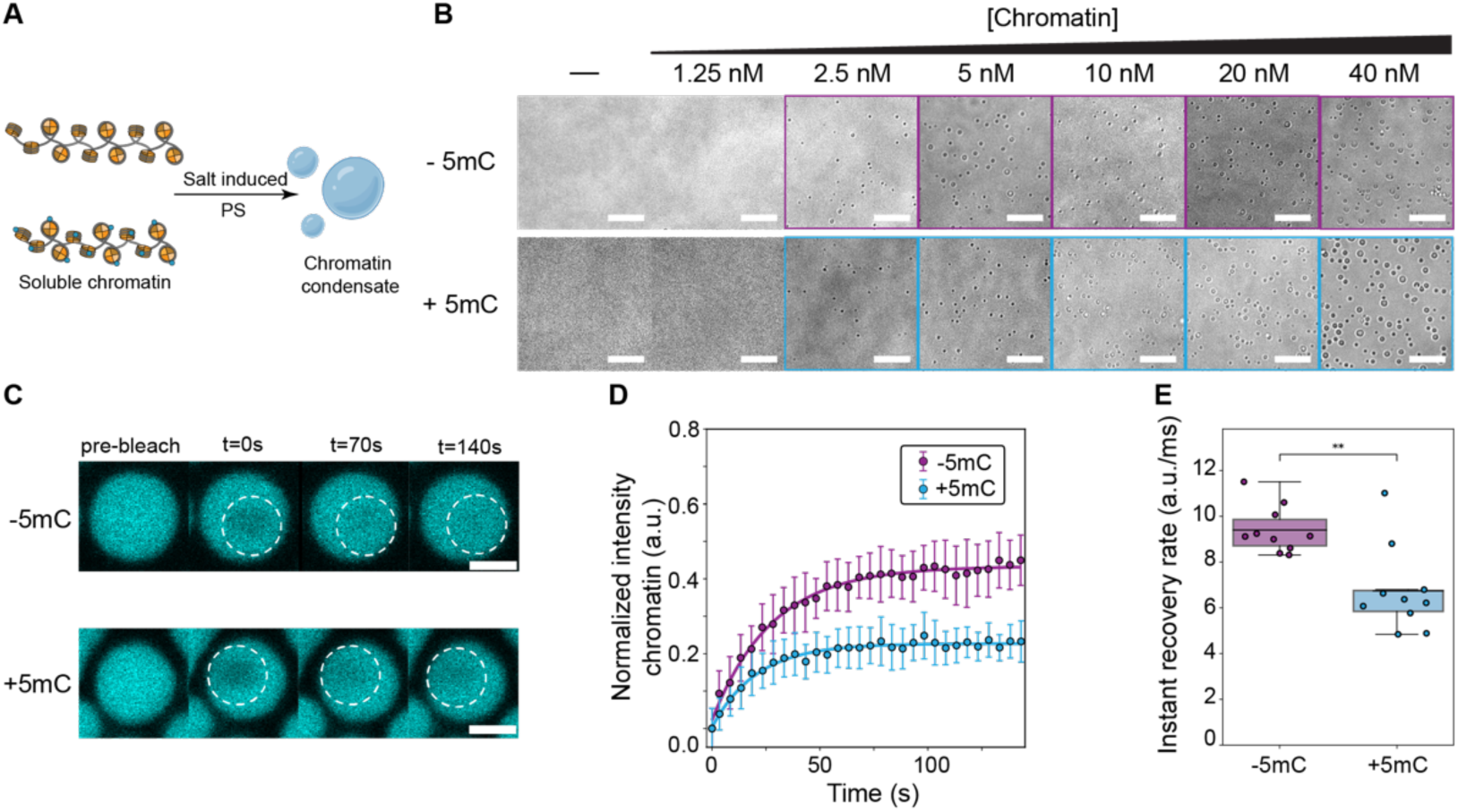
CpG methylation modulates chromatin condensate dynamics. **A)** Schematic of the chromatin phase separation assay induced by salt. **B**) Increasing concentrations of −5mC and +5mC chromatin fibers were incubated at 150 mM NaCl and imaged by bright-field optical microscopy. Squares highlight conditions in which condensates are observed. Scale bar = 10 µm. **C**) Representative microscopy images of Cy5-H2B labeled chromatin droplets before and after partial photobleaching. Scale bar = 2 µm **D**) Normalized fluorescence recovery curves for –5mC and +5mC condensates (n = 10 droplets per condition). Mean is reported and error bars represent standard deviation. **E**) Instantaneous recovery rates, derived from the initial slopes of the recovery curves, show slower chromatin dynamics in +5mC condensates compared to −5mC.

Given the observed differences in nucleosome wrapping and fiber architecture (**Fig. 2**), we next asked whether CpG methylation affects chromatin dynamics within phase-separated assemblies, even when the threshold concentration for phase separation remains unchanged. To test this, we incorporated Cy3-labeled H2B histones into chromatin fibers and performed fluorescence recovery after photobleaching (FRAP). A defined region within each condensate was photobleached, and fluorescence recovery was monitored over time (**Fig. 3C**). Remarkably, both the extent (**Fig. 3D**) and rate (**Fig. 3E**) of recovery were reduced in +5mC chromatin compared to -5mC, indicating that both the diffusion rate and chromatin mobile fraction are reduced within +5mC condensates.

These results indicate that CpG methylation restricts chromatin mobility, potentially by altering inter-nucleosome interactions and the internal connectivity of the condensate network.

### CpG Methylation Produces Denser and More Viscoelastic Chromatin Condensates

The reduced mobility of +5mC chromatin within condensates suggested that CpG methylation may also alter the condensates’ underlying material properties. To directly probe their viscoelastic behavior, we performed video particle tracking (VPT) nanorheology^46, 47^ (**Fig. 4A**). In this approach, the motion of 200 nm fluorescent beads, which were passively embedded into the chromatin condensates, was tracked over time. Beads were more dynamic in -5mC than in +5mC chromatin condensates, as evident from the spatial extent of the individual bead motion (**Fig. 4B**) and significantly higher ensemble-average mean-squared displacement (MSD) of beads in the unmethylated condition (**Fig. 4C**). In both samples, the beads showed subdiffusive motion with the diffusivity parameter, 𝛼 <1, signifying the viscoelastic nature of the condensates arising from a strong intra-condensate nucleosome interaction network.

**Figure 4.**
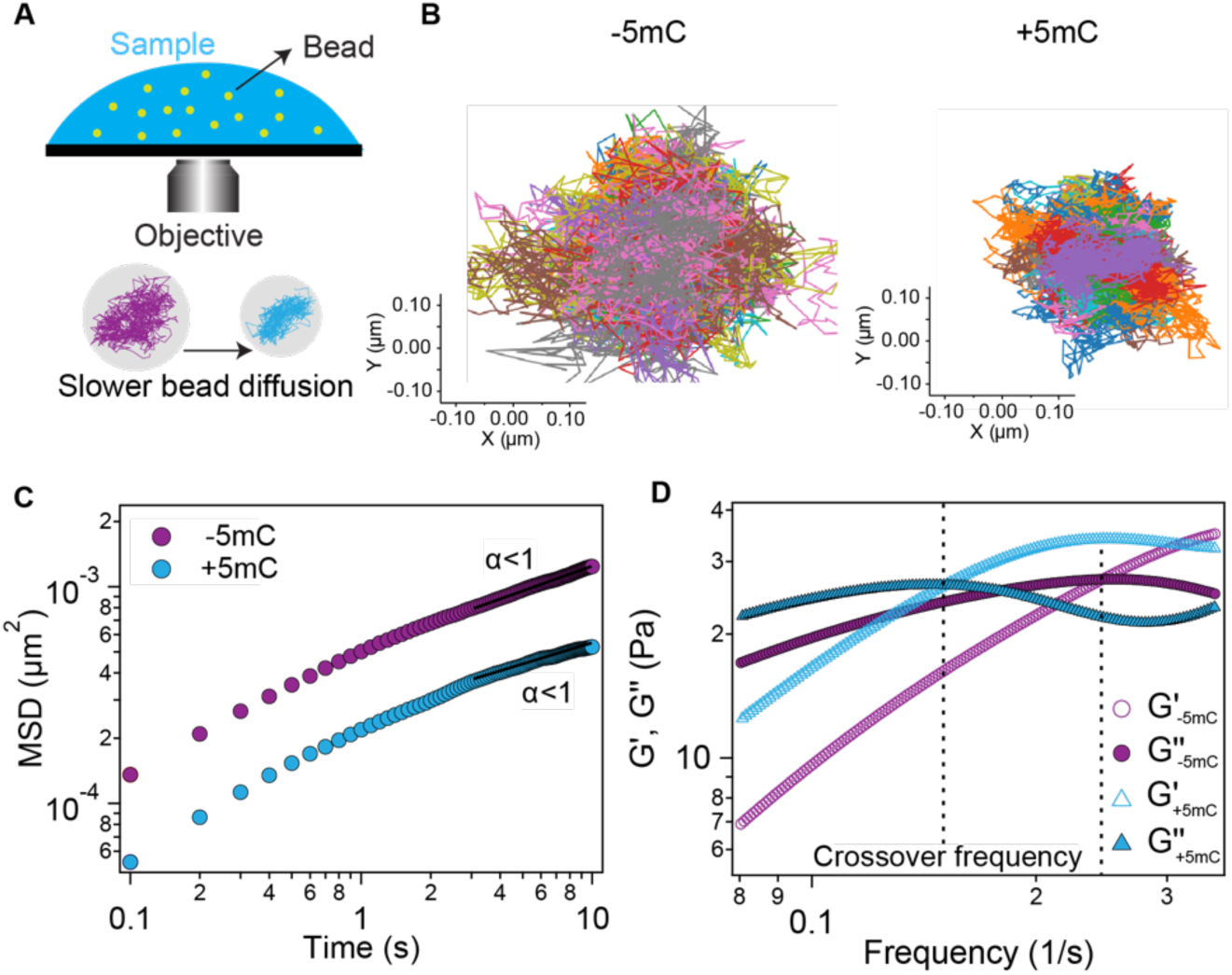
CpG Methylation Alters the Material Properties of Chromatin Condensates. **A)** Schematic representation of VPT nanorheology employed to quantify the material properties of chromatin condensates. The upper panel shows beads embedded inside a condensate and the lower panel depicts bead trajectories with altered diffusivity inside condensates with different material properties. **B**) Trajectories of the beads tracked inside the -5mC (left) and +5mC (right) chromatin condensates. Different colors represent trajectories of distinct beads tracked inside the condensates. **C**) Ensemble-averaged mean-squared displacement (MSD) of 200 nm carboxylate-modified beads inside -5mC and +5mC chromatin condensates. The beads exhibited subdiffusive motion inside both type of condensates, depicted by the diffusivity parameter, 𝛼<1. **D**) The storage and loss moduli of the -5mC and +5mC chromatin condensates were estimated utilizing the Evans method^48^ from the MSDs shown in C. The dotted lines depict the frequency at which the crossover between storage modulus (Gʹ) and loss modulus (Gʹʹ) was observed for these condensates.

To quantify viscoelasticity, we employed Evans method and utilized MSDs estimated from VPT measurements as input to it to compute the complex shear moduli^46, 48^(**Fig. 4D**). The frequency-dependent viscoelastic moduli revealed that both types of chromatin condensates (±5mC) exhibit characteristic viscoelastic behavior. Furthermore, +5mC chromatin condensates showed a crossover between the storage modulus (Gʹ) and the loss modulus (Gʹʹ) at a lower frequency compared to -5mC chromatin condensates. The inverse of this frequency, termed crossover frequency, provides an estimate of the terminal relaxation time (𝜏) of the interaction network, which reflects the timescale required for the condensate network to reorganize. +5mC condensates exhibited a longer relaxation time (𝜏_+msc_∼6.67 ± 0.56 s) compared to -5mC chromatin condensates (𝜏_-msc_∼4.02 ± 0.95 s), indicating CpG methylation produces a more viscoelastic chromatin network.

Because these mechanical differences suggested corresponding changes in condensate architecture in the nanoscale, we next examined their nanoscale organization using cryo-electron tomography (cryo-ET). Chromatin condensates formed with 25-bp linker arrays at physiological salt were vitrified by plunge freezing, imaged with tilt series acquisition, and reconstructed into tomograms that were denoised to enhance structural features. The tomograms showed subtle qualitative differences in the internal organization of -5mC versus +5mC chromatin condensates (**Fig. S4A, B**). −5mC condensates displayed a more open and porous architecture, with more visible unoccupied volumes, whereas +5mC condensates presented a more densely packed internal organization with reduced void space. These ultrastructural differences are consistent with the increased viscoelastic properties of CpG-methylated condensates measured by VPT (**Fig. 4C, D**).

Together, these observations indicate that CpG methylation does not substantially alter the threshold concentration for chromatin phase separation but instead modulates the material properties and nanoscale network structure of the resulting condensates. Rather than altering the overall thermodynamic driving force for condensation or the multivalent interactions among nucleosomes, CpG methylation reorganizes the microscopic architecture of the condensate, producing a denser and more crosslinked chromatin network. This altered ultrastructure is reflected in slower relaxation dynamics and increased viscoelasticity, suggesting that CpG methylation tunes the mechanical and dynamic behavior of higher-order chromatin assemblies through changes in DNA mechanics and inter-nucleosome stacking.

### CpG Methylation Promotes HP1α-Chromatin Interactions via Chromatin Fiber Properties

Chromatin compaction and gene silencing within heterochromatin are orchestrated by an interplay between DNA methylation and the heterochromatin protein 1α (HP1α). HP1α recognizes the histone mark H3K9me3 through its chromodomain and oligomerizes to bridge nucleosomes, thereby promoting chromatin condensation^21,22,49^. HP1α is a key component of repressive chromatin compartments and can itself drive chromatin phase separation *in vitro*^18,23^. Notably, H3K9me3 and CpG methylation frequently co-occur within constitutive heterochromatin, where they are thought to act synergistically to stabilize compact, transcriptionally silent chromatin domains^50–52^. This close spatial and functional association raises the possibility that CpG methylation influences how HP1α interacts with and organizes chromatin.

To test this, we reconstituted chromatin fibers containing H3K9me mimics generated by methyl-lysine analog chemistry^53^ in the presence or absence of CpG methylation and measured the concentration of HP1α required to induce condensate formation (**Fig. 5A**). At a fixed chromatin concentration (20 nM), -5mC chromatin required four-fold more HP1α to phase separate than +5mC chromatin (5 μM versus1.25 μM) (**Fig. 5B**). This reduction in threshold concentration indicates that CpG methylation enhances HP1α–chromatin interactions and facilitates condensation.

**Figure 5.**
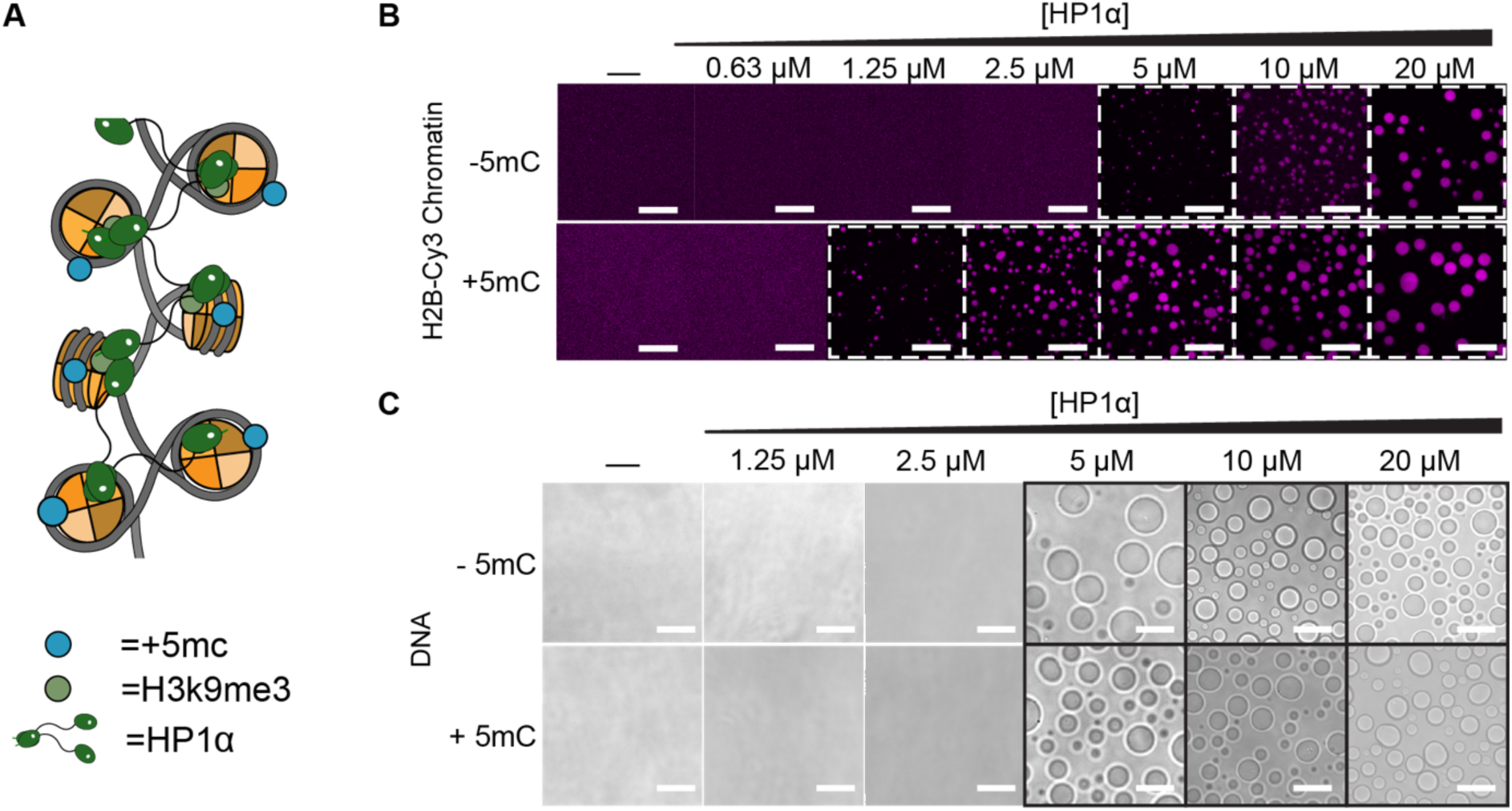
CpG Methylation Modulates HP1α-Chromatin Interactions. **A)** Schematic illustrating interactions between HP1α and chromatin fibers carrying H3K9me3 and CpG methylation. **B**) Confocal microscopy images of the indicated chromatin condensates formed at increasing HP1α concentrations. Chromatin is fluorescently labeled. White dashed squares indicate conditions in which condensates are observed. Scale bar = 20 µm. **C**) Bright-field images of condensates formed by HP1α and dsDNA. Fixed concentrations of -5mC and +5mC dsDNA were incubated with titrated HP1α. Black squares mark conditions in which condensates are observed, showing comparable threshold concentrations for −5mC and +5mC DNA. Scale bar = 20 µm.

We next asked whether this enhancement reflects higher affinity for CpG methylated DNA by HP1α or instead arises from changes in chromatin fiber properties that indirectly affect the accessibility or valency of HP1α binding sites. To distinguish between these mechanisms, we assessed HP1α phase separation with naked dsDNA (2 kb) with or without CpG methylation. Both substrates displayed identical HP1α threshold concentration, indicating that CpG methylation does not alter HP1α binding through direct DNA contacts (**Fig. 5C**). Thus, the increased propensity of +5mC chromatin to undergo HP1α-driven phase separation is not due to HP1α recognizing methylated DNA per se, but rather to methylation-dependent biophysical changes within the chromatin fiber that promote a more favorable HP1α engagement.

Together, these results indicate that CpG methylation stabilizes HP1α–chromatin interactions not by altering HP1α binding to naked DNA, but indirectly through changes in the chromatin viscoelastic network that enhance local chromatin stability and HP1α binding.

## Discussion

DNA methylation is a central epigenetic modification that contributes to transcriptional silencing and heterochromatin formation, yet the physical mechanisms by which it influences chromatin architecture remain incompletely understood. Here, by combining single-molecule force spectroscopy, phase separation assays, nano-rheology, and cryo-ET, we show that CpG methylation modulates the physical properties of chromatin across multiple scales, from naked dsDNA and individual nucleosomes to chromatin fibers and condensates, and finally to their interactions with the heterochromatin protein HP1α.

At the level of DNA mechanics, CpG methylation increases the contour and persistence length (**Fig. 1**), indicating a slight extension of the helical structure and reduced bending flexibility. These observations are consistent with previous biophysical studies reporting that methylation increases DNA contour length^54^ and stiffness^29,35–39^. However, previous work also reported a decreased persistence length for methylated DNA^54^, underscoring that methylation effects can be sensitive to sequence context, methylation density, and ionic conditions^55^. Within our experimental framework, the modest increase in stiffness provides a physical basis for how methylated DNA might engage differently with the histone octamer.

Indeed, at the nucleosome level we find that CpG methylation affects DNA unwrapping. Single-molecule measurements reveal lower forces for inner turn unwrapping and reduced energy required to unwrap the outer turn (**Fig. 2**). The latter is at least in part due to altered DNA stiffness. This combination suggests that methylated DNA, while mechanically more rigid, wraps less stably around the octamer, likely resulting in subtle changes in DNA geometry and steric packing at the nucleosome periphery. The resulting decrease in wrapping stability could facilitate additional nucleosome loading and compaction, in line with earlier theoretical predictions suggesting that greater DNA stiffness leads to increased nucleosome formation^56^. These findings support a model in which CpG methylation promotes local compaction of individual fibers while simultaneously weakening individual octamer–DNA contacts and enhancing DNA breathing motions^57^. This dual effect highlights how DNA methylation reshapes the energy landscape of chromatin by coupling altered DNA mechanics to nucleosome stability and wrapping geometry.

At the level of high-order chromatin organization, salt-driven phase separation assays revealed that CpG methylated and unmethylated arrays exhibit similar threshold concentrations for condensate formation (**Fig. 3B**), indicating that CpG methylation does not strongly alter the overall thermodynamic driving force or the effective valency of multivalent inter–nucleosome interactions required for phase separation. However, within condensates, FRAP revealed that methylated chromatin is less mobile (**Fig. 3C-E**): both the rate and extent of recovery are reduced, consistent with slower diffusion and a smaller mobile fraction of chromatin in the dense phase. Thus, while the onset of phase separation is largely unchanged, the internal dynamics of the condensed state are substantially altered and stabilized by CpG methylation. Video particle tracking nano-rheology further revealed that CpG methylation slows the relaxation of the condensate network and increases its viscoelasticity (**Fig. 4**). Beads diffusing within methylated condensates display lower MSDs and longer terminal relaxation times than those in unmethylated condensates, indicating that the chromatin network in the dense phase is more resistant to rearrangement. Together with the Cryo-ET images (**Fig. S4**), these orthogonal measurements indicate that CpG methylation does not change whether chromatin phase-separates but instead remodels the microstructure and material properties of the condensates once formed, producing a tighter, more crosslinked network with slower internal dynamics.

Finally, we find that CpG methylation enhances the responsiveness of chromatin to HP1α, a key effector of heterochromatin organization (**Fig. 5**). CpG methylated chromatin requires a lower concentration of HP1α to undergo phase separation. Importantly, this effect is not observed when HP1α is tested on naked dsDNA, indicating that CpG methylation does not act by directly modulating HP1α–DNA affinity. Instead, CpG methylation appears to act indirectly by reorganizing chromatin architecture and generating a denser condensate network that provides more favorable or persistent HP1α bridging configurations. In this way, CpG methylation can potentiate HP1α-mediated compaction and stabilization of heterochromatin without dramatically shifting the bulk condensation threshold.

Taken together, our results support a multiscale model in which CpG methylation tunes chromatin mechanics and dynamics rather than acting solely as a static binding platform for reader proteins. At the molecular level, methylation stiffens DNA and weakens nucleosomal wrapping; at the condensate level, it produces denser and more viscoelastic networks; and at the level of effector engagement, it enhances HP1α-driven condensation. By reshaping the physical landscape of chromatin across these scales, DNA methylation biases chromatin toward stable, low-mobility configurations characteristic of repressive, heterochromatic states. More broadly, our findings illustrate how epigenetic modifications can modulate genome function not only through biochemical signaling pathways but also by tuning the physical and material properties of chromatin itself.

## Materials and Methods

### Cloning

Restriction-ligation cloning was used to generate a 12×601 NRL172 DNA construct for single-molecule force-spectroscopy. The vector pSuperCos and the 12×601 NRL172 donor plasmid pWM_12×601_25bpLinker (Addgene, plasmid #157788) were digested with BamHI and DrdI (5 U/μg). Fragments were gel-purified (0.8% agarose in 1xTAE, 0.5 μg/ml ethidium bromide) using the NucleoSpin Gel & PCR Clean-up kit (MACHEREY-NAGEL) and ligated with T4 DNA ligase (ThermoFisher). Ligation mixtures were transformed into chemically competent DH5α and plasmid DNA of single colonies was isolated using the Wizard® Plus SV Minipreps DNA Purification System (Promega). Plasmid sequencing was performed by Plasmidsaurus using Oxford Nanopore Technology.

### Optical Tweezer Construct Preparation

12×601 NRL172 plasmid DNA for single-molecule experiments was amplified in methyltransferase deficient (*dam^-^/dcm^-^*) chemically competent *E. coli* (NEB, C2925). Plasmid DNA was linearized with NotI (5 U/μg) yielding a construct with DNA handles of approximately 2.7 and 2.0 kb flanking the 601 arrays. 5’oOverhangs were filled with dGTP and biotin-16-dCTP using Klenow fragment (Jena Bioscience). Biotinylated DNA was methylated using the CpG methyltransferase M.SssI (NEB). The degree of methylation was assessed by digestion using the methylation-sensitive restriction enzyme HpaII and digestion patterns were assessed with gel electrophoresis (1.2% agarose in 1xTAE, 0.5 μg/ml ethidium bromide). The used DNA construct has a GC content of 51% and contains 482 GC dinucleotides. All DNA purifications were performed with the NucleoSpin Gel & PCR Clean-up (MACHEREY-NAGEL).

### Chromatin Reconstitution

#### Eeftens lab

Chromatin reconstitution was performed by salt gradient dialysis in 7K MWCO Slide-A-Lyzer MINI Dialysis Devices (ThermoFisher) using a multi-headed peristaltic pump (Masterflex) based on previously established protocols^58^. Reconstitution mixtures contained 0.179 M 601 DNA in addition to recombinant human histone octamers (EpiCypher, 16-0001) with increasing molar ratios of histone octamer to 601 DNA. An electrophoretic mobility shift assay (EMSA) was conducted to determine the degree of saturation of nucleosome assembly at the 601 sequences. To this end, reconstituted chromatin samples were digested with NlaIII (10 U/μg) and analyzed using gel electrophoresis (0.8% agarose in 0.2x TB, 0.5 μg/ml ethidium bromide). Nucleosomal arrays were stored at 4°C for up to 6 weeks to ensure consistent sample quality.

#### Sanulli lab

All human histones were expressed and purified from *E. coli* following published protocols^59^. Methyl lysine analogue (MLA) containing H3 histones at position 9 (H3Kc9me3) were prepared as described previously^60^. Histone labeling was performed following previously described protocol^61^. Histone octamers were assembled from purified histones by salt dialysis as described^23^ and purified by size exclusion using a Superdex-200 column. DNA was generated by restriction enzyme digestion of a plasmid containing 12 consecutive 601s spaced by 25 bp, followed by sizing exclusion purification, as previously described^23^. The reconstitution of nucleosome arrays followed the protocols described^59^. Histone octamers are combined in equimolar amount with 12-mer DNA (12 repeats of the 601 DNA sequence separated by 25-bp linkers). Final dialysis against TCS buffer (20 mM HEPES pH 7.5, 0.1 mM EDTA) was performed overnight. Assembled arrays were used for experiments within 5 days after assembly. Since the 601 repeats in the 12-mer DNA sequence are separated by BstXI restriction enzyme sites, BstXI digestion was performed followed by native gel analysis. Over-assembly was avoided by ensuring that >95% of the digested fragments migrated as mononucleosomes rather than slower migrating species. Fluorescently labeled chromatin was prepared by incorporating 5% of octamers containing Cy3 or Cy5-labeled H2B^61^.

### 12×601 DNA Methylation

Purified 12×601 DNA was incubated with M.SssI CpG methyltransferase (NEB #M0226M) in NEBuffer rCutSmart (NEB) supplemented with S-adenosylmethionine (SAM #B9003S) according to the manufacturer’s recommendations. Methylation reactions were carried out at 37°C for 4 h with gentle mixing to ensure uniform accessibility of CpG sites. Following incubation, reactions were terminated by placing on ice and cooling to 4°C. Successful methylation was confirmed by the expected protection of BsiWI-sensitive site, which selectively cleaves unmethylated CpG sites. Methylated DNA was purified by ethanol precipitation and sequential 70% ethanol washes to remove residual methyltransferase, buffer components, and cofactors. The DNA pellet was briefly air-dried and resuspended for downstream chromatin assembly.

### Single-Molecule Force Spectroscopy

Force spectroscopy experiments were performed with dual optical traps in a multi-channel laminar flow cell connected to a microfluidics system (C-Trap, LUMICKS). Prior to all measurements, the flow cell was cleaned and subsequently passivated using 20 mM HEPES pH 7.5, 150 mM NaCl, 0.5% (w/v) casein and 0.01% (w/v) BSA. Measurements were performed in a buffer containing 20 mM HEPES pH 7.5, 100 mM NaCl, 2 mM MgCl_2_, 0.2% (w/v) BSA and 0.02% (v/v) Tween 20. To perform force spectroscopy measurements, two streptavidin-functionalized polystyrene beads with a diameter of 1.76 µm (Spherotech BV) were optically trapped and a force calibration was conducted. Biotinylated DNA or chromatin was flow-stretched and tethered between the trapped beads. Prior to each measurement, the force offset was set to 0 pN. Force spectroscopy measurements were performed with a constant trap velocity of 20 nm/s and forces on the stationary bead were registered using back focal plane detection of scattered optical trapping light.

### Data Analysis and Statistics

Data analysis was performed in Python (version 3.9.7) with custom-written scripts using the lumicks.pylake package (version 1.2.0). Force trace data between 1 and 30 pN for DNA was fitted to Odijk’s extensible worm-like chain (eWLC) with force as the dependent variable to robustly solve L*_p_* and L*_c_*^34,62,63^. Determination of the area under the curve was achieved by numerical integration (SciPy Simpson). All F,d-curves were included to determine inner turn unwrapping forces. Unwrapping forces were plotted for all nucleosomes in case of undersaturated or saturated fibers. The first 12 unwrapping events were included for oversaturated fibers. To obtain average F,d-curves in the low force regime, data was interpolated and averaged. Data was tested for normality using the Shapiro-Wilk test. Normally distributed datasets were compared using Student’s t-tests and non-normally distributed datasets were compared with the Mann-Whitney U-test. In all cases, the threshold of significance was set at p < 0.05.

### hHP1α Purification

N-terminally 6X-His tagged hHP1α proteins (WT and Hinge mutants) were purified from *E. coli* as previously described^23^. Briefly, hHP1α was affinity purified with His-Trap followed by TEV protease treatment to cleave the N-terminal 6x-His tag. After TEV cleavage, it was subjected to HiTrap Q HP column (GE Healthcare) and Superdex 200HR 10/300 column (GE Healthcare). hHP1α is stored in 25 mM HEPES pH 7.5, 150 mM KCl, 1 mM DTT and 10% glycerol. Concentrations were measured by UV absorbance at 280 nm and using the calculated extinction coefficient ε=29,495.

### Phase Separation Assay

Salt induced chromatin condensates were generated by mixing 1:1 ratio 2X chromatin stock stored in TCS buffer and 2X phase separation buffer (300 mM NaCl, 20 mM HEPES pH 7.5, 5% glycerol), to achieve final 150 mM NaCl. Phase separation assays with HP1α were performed by mixing 2X chromatin stock stored in TCS buffer and HP1α stored in 150 mM NaCl, 20 mM HEPES pH 7.5, 5% glycerol at 1:1 ratio to achieve the indicated concentrations. All samples were immediately loaded onto 384-well glass-bottom plates (Greiner, #781892) precoated with PEG-silane (Laysan Bio MPEG-SIL) as previously described^23,64^ to minimize nonspecific surface interactions. Droplets were imaged using a Nikon Eclipse Ts2 microscope equipped with a 40X air objective within 2 hours. Image contrast and brightness were uniformly adjusted using Fiji (ImageJ). Confocal fluorescence images were acquired on either a Zeiss LSM 880 microscope (40X oil objective) or a Nikon Spinning Disk Confocal Microscope (60X oil objective). Acquisition settings were kept constant within each experimental series.

### Fluorescence Recovery After Photobleaching (FRAP)

FRAP experiments were performed by photobleaching Cy3-labeled proteins within condensates using 100% laser intensity at 594 nm. For chromatin condensates, H2B-Cy3 and HP1α-Cy3 were selectively bleached, and fluorescence recovery was monitored over time using time-lapse imaging at low laser power to minimize secondary bleaching. H2B-Cy3 was labeled with 50% labeling efficiency quantified spectroscopically, and the final loading of labeled biomolecules was 5%. To minimize droplet movement, BSA was omitted in the glass passivation protocol for FRAP experiments. Each recovery trace was background-subtracted and normalized to pre-bleach intensity. Experiments were performed in triplicate with at least 10 droplets analyzed per condition.

### FRAP Data Analysis

Quantitative analysis of FRAP data was performed using custom Python scripts using the data provided by the Zeiss software upon acquisition. Time–intensity traces were shifted for each droplet and aligned such that *t = 0* corresponded to the midpoint of the bleach pulse. Each trace was background-subtracted, normalized to its pre-bleach intensity, and baseline-corrected to account for slow photobleaching during recovery. Normalized traces from at least ten droplets per condition were averaged to obtain mean recovery profiles. The early recovery phase was fit with a linear model to determine the instantaneous recovery rate, which reflects the initial diffusion-limited dynamics within condensates. Statistical comparisons between conditions (e.g., −5mC vs. +5mC) were performed using two-tailed unpaired *t*-tests, with significance reported according to standard conventions (*p* < 0.05 = *, *p* < 0.01 = **, *p* < 0.001 = ***, *p* < 0.0001 = ****).

### Electron Microscopy

Cryo-ET samples were prepared on Quantifoil Cu 1.2/1.3 200-mesh grids (Electron Microscopy Sciences) that were glow-discharged for 90 s at 10 mA using a PELCO easiGlow system prior to sample application. Chromatin condensate solutions (3 µL) were applied to the grids, and vitrification was performed using a Mark IV Vitrobot (Thermo Fisher Scientific) with a blot time of 1 s, blot force of 1, at 4°C and 100% relative humidity. Grids were plunge-frozen into liquid ethane and stored in liquid nitrogen until screening and data acquisition.

### Cryo-ET Data Acquisition and Processing

Grids were imaged with a Titan Krios G4 at 300 kV equipped with an X-FEG electron gun. Data were acquired with a Falcon 4i direct electron detector and SelectrisX energy filter set to a 10 eV slit width. Tilt series were acquired using Tomo 5 (Thermo Fisher Scientific) and a 70 µm objective lens aperture. The total dose was ∼180 e/Å^2^ linearly split over 33 tilts with 3° increments and tilt range of -48° to 48°. A pixel size of 1.2 Å/px and nominal target defocus range of 2-5 µm was used for acquisition. A total of 143 and 138 tilt series were acquired for the unmethylated and methylated conditions, respectively.Tilt series were pre-processed on-the-fly using AreTomoLive^65^. This includes gain, motion and CTF correction, followed by patch-based tilt series alignment and tomogram reconstruction with weighted back-projection. Tomogram thickness estimates were obtained from the AreTomoLive pre-processing metrics. Tomograms were denoised with DenoiseET^1^ using a pre-trained model from tomograms of synaptosomes (Cryo-ET Data Portal ID DS-10443). Search map montages and representative slices of denoised tomograms were visualized using IMOD^66^.

### Sample Preparation for Nano-rheology

Chromatin condensates were prepared at a chromatin concentration of 400 ng/µL in the buffer containing150 mM NaCl, 10 mM HEPES pH 7.5, 2.5% glycerol. The prepared samples, in a 10 µL volume, were incubated for 30 minutes in a tube and then transferred to an 18 x 18 mm microscope coverslip coated with Sigmacote (Sigma Aldrich) solution. The coverslips were coated by immersing them in Sigmacote solution for ∼2 minutes. After 2 minutes, the coverslips were subsequently dried using compressed air and then incubated at 37°C overnight. The prepared samples were sandwiched between a 75 x 25 x 1 mm thick microscope glass slide using five layers of double-sided tape (Scotch 3M). Further, to prevent evaporation, ∼200 µL of mineral oil was injected into the chamber to surround the sample.

### Video Particle Tracking (VPT) Nanorheology

The samples for measurements were prepared as described in the previous section, with an additional step of carboxylate-modified yellow-green fluorescent 200 nm microspheres (Invitrogen) added to the sample before the induction of phase separation. The samples were then moved on to a Zeiss Primovert inverted microscope with a 100x oil-immersion objective lens and equipped with a Teledyne FLIR Blackfly S USB3 CMOS camera. The motion of the beads inside the chromatin condensates was monitored by acquiring a 1000-frame video at a rate of 10 frames per second with a camera exposure time of 100 ms. All the measurements were repeated over 3 independently prepared samples.

### VPT Data Analysis

The VPT nanorheology framework is described in detail in our previous work^46,47^. In brief, the TrackMate plugin of the ImageJ FIJI software was used for tracking the bead’s motion inside the condensates^67^. The tracking provided 2D trajectories of the beads, which were further analyzed. For correcting any possible drift during the measurements, the diffusion of the center of mass of the beads was estimated from the center of mass vector 𝑹.

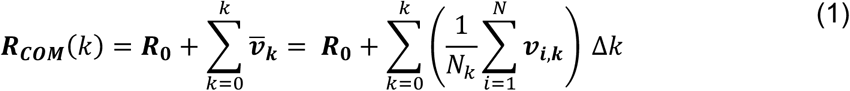

Here, 𝑹_𝟎_ is the initial center of mass vector calculated by averaging the coordinates of all beads in the first frame (𝑘=1), 𝒗-_𝒌_ is the mean velocity of all particles in the frame 𝑘, 𝒗_𝒊,𝒌_ is the velocity of the particle 𝑖 in frame 𝑘 in units of µm/frame and 𝑁_+_ is the number of particles in the frame 𝑘. The quantity Δ𝑘 is the frame difference, which is 1 in the present case. The drift correction was done by subtracting the center of mass from the individual bead trajectories. This was used to calculate the ensemble-averaged mean-squared displacement (MSD) using:

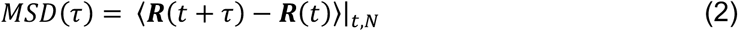

In Eq. (6), 𝜏 is the lag time. To understand the nature of the bead diffusion inside the chromatin condensates, we fitted the MSD to the generalized Stokes–Einstein relation,

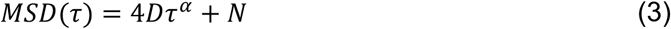

Here, 𝐷 is the diffusion coefficient, 𝛼 is the diffusivity exponent, and 𝑁 is a term to account for the tracking noise.

### Estimation of dynamical moduli from nanorheology measurements

The complex and the loss moduli for the -5mC and +5mC chromatin condensates were estimated using the Evans method^48^ (a custom Python script was used). The details of the complex moduli estimation from MSD are described elsewhere^68^. Briefly, MSD estimated from the VPT nanorheology measurements was used to obtain the compliance 𝑗(𝑡) using equation:

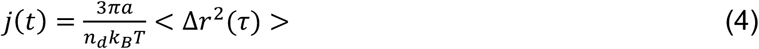

Here, 𝑎 represents the probe particle radius (𝑎=100 nm), 𝑛_d_ = 2 represents the dimension of the probe particle trajectories based on which the MSDs were estimated, 𝑘_<_ represents the Boltzmann constant, and 𝑇 represents the absolute temperature at which the rheology measurements were done.

The complex shear modulus and the compliance are related through a convolution integral

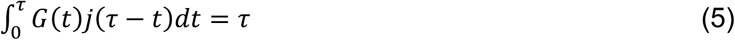

This provides an opportunity to compute complex shear modulus by performing a Fourier transform using the relation,

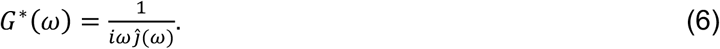

We further followed the methodology of Evans, R. et al.^48^ to estimate the frequency-space complex moduli 𝐺^∗^(𝜔) using the discrete experimental data points (𝑡_1_, 𝑗_1_) using the relation,

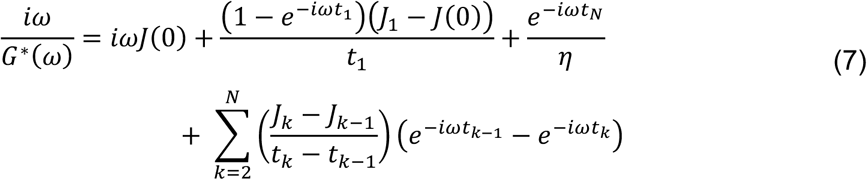

Where 𝑗(0) is acquired through the extrapolation of the experimental data for compliance 𝑡 = 0 using a linear fit of the initial four data points and viscosity, 𝜂 using a linear fit of the final ten data points following the relation, 𝜂 = 1/𝚥(𝑡).

## Acknowledgements

We thank members of the Sanulli laboratory for helpful discussions and suggestions. Work in the Sanulli laboratory was supported by the Chan Zuckerberg Biohub (to SS), NIH DP2GM149752 (to SS), the Searle Scholars Program (to SS), and Propel postdoctoral fellowship (to AR). The Eeftens lab thanks Tieke Kuijpers and Kes van Blitterswijk for early contributions to the project. Work in the Banerjee laboratory was supported by National Institutes of Health grant R35 GM138186 and St. Jude Research Collaborative on Biophysics of RNP granules.

## Author information

These authors contributed equally: Daris LJC, Rizvi A.

## Contributions

AR, LJCD, JME and SS conceived the project and wrote the manuscript. LJCD performed the single molecule experiments, with methodology by LAL. AR performed phase separation and FRAP assays. AS performed the VPT experiments and analyzed the data with support from PRB. AR prepared EM samples, and JH, EAM and DK processed the data. All authors discussed the results and commented on the manuscript.

## Ethics declarations

The authors declare no competing interests.

## Supplementary Figures

**Figure S1.**
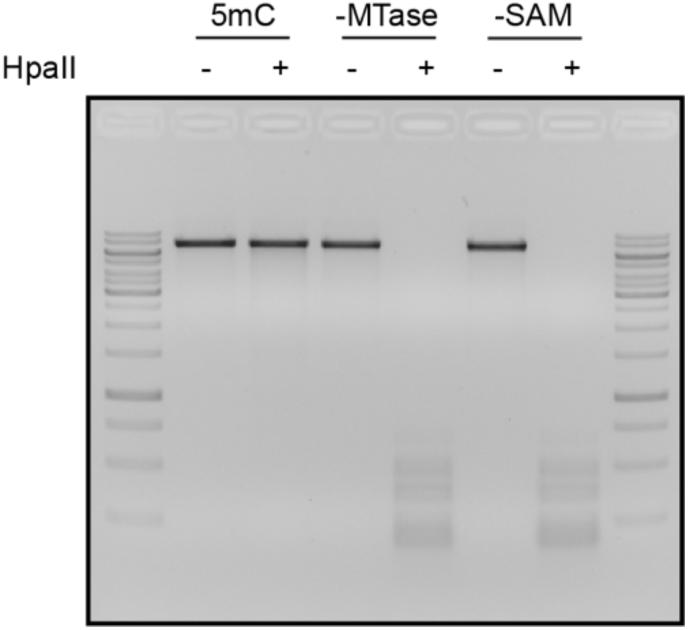
Verification of CpG methylation of the DNA Construct. The CpG methyltransferase (MTase) M.SssI was used to methylate all cytosines at CpG sites withing the DNA construct. Digestion with the methylation-sensitive restriction enzyme HpaII confirmed complete methylation: the DNA was protected from cleavage following MTase treatment (lane 3). In the absence of MTase or SAM, the DNA remained sensitive to HpaII digestion (lanes 5 and 7). The fully methylated DNA construct was used for force spectroscopy and as the template for chromatin reconstitution.

**Figure S2.**
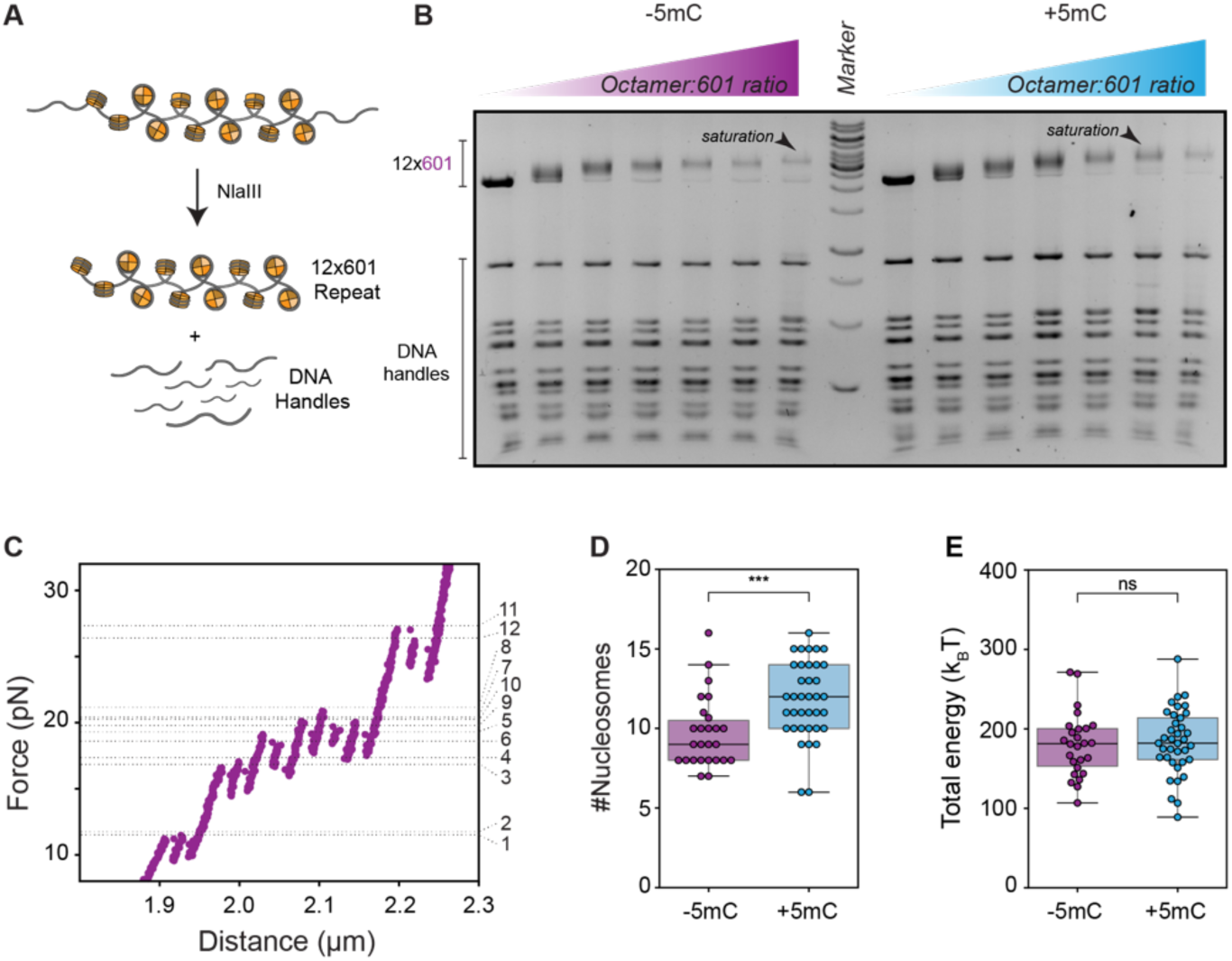
Determining the Number of Nucleosomes per Chromatin Fiber. **A)** Enzymatic validation of nucleosome assembly. Reconstituted chromatin fibers were digested with NlaIII to distinguish between nucleosome assembly at the 601 sequences from the assembly on the DNA handles. **B**) Saturation of nucleosome assembly determined by EMSA. Chromatin fibers were reconstituted with increasing octamer:DNA molar ratios to ensure full saturation of the 601 sites. NlaIII digestion releases the 12×601 segment (upper bands) and further digested the DNA handles (residual lower bands). Progressive mobility shifts of the 601 reflects increasing nucleosome occupancy until saturation, whereas excess octamer results in additional shifts corresponding to over-assembly on the DNA handles. **C**) Discrete unwrapping steps in the high-force regime correspond to inner turn rupture events, enabling quantification of nucleosome number per tether. Peak forces before each rupture event were used to identify individual unwrapping transitions. **D**) Quantification of ruptures events shows a significant higher number of nucleosomes per chromatin fiber in the +5mC condition (Mann-Whitney U-test: p < 0.001, n = 37 and n = 26 for +5mC and -5mC, respectively). **E**) Total stretching energy (area under the curve in the low-force regime) is not significantly different between -5mC and +5mC fibers (Student’s t-test: ns, n = 37 and n = 26 for +5mC and -5mC, respectively). This apparent similarity results from increased nucleosome occupancy in the 5mC condition, rather than unchanged fiber mechanics.

**Figure S3.**
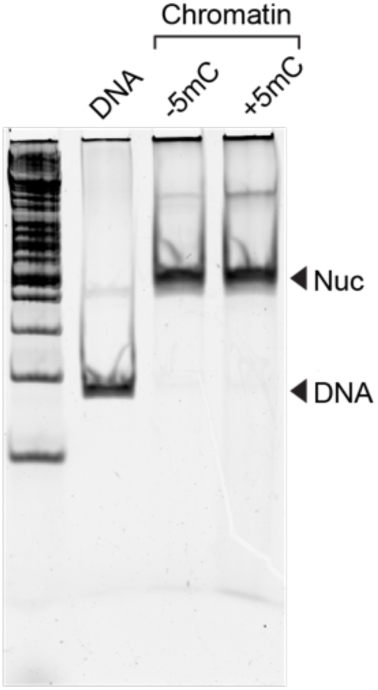
Chromatin Quality Control. Enzymatic validation of nucleosome assembly. Naked dsDNA and reconstituted chromatin fibers were digested with BstXI and analyzed on a native gel. Both -5mC and +5mC chromatin show complete nucleosome occupancy.

**Figure S4.**
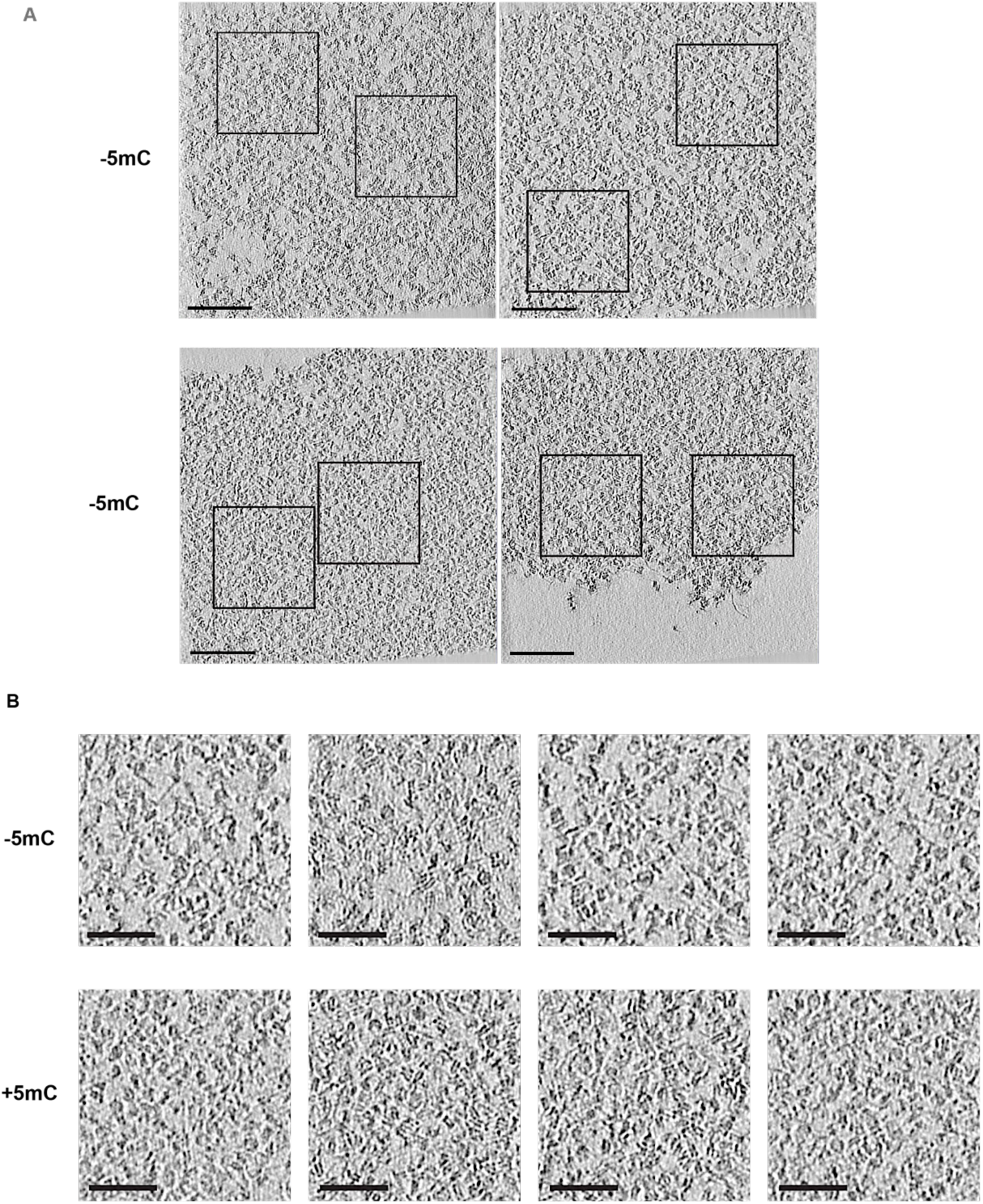
Qualitative assessment of chromatin condensates by Cryo-ET. **A)** Representative enhanced views generated by summing five consecutive tomogram slices of -5mC (top) and +5mC (bottom) chromatin condensate. **B)** Magnified views of the regions indicated by black squares in A, highlighting differences in internal organization. Scale bar = 50 nm

